# Pervasive Adaptation of Hepatitis C Virus to Interferon Lambda Polymorphism Across Multiple Genotypes

**DOI:** 10.1101/484071

**Authors:** Nimisha Chaturvedi, Evguenia S. Svarovskaia, Hongmei Mo, Anu O. Osinusi, Diana M. Brainard, G Mani Subramanian, John G. McHutchison, Stefan Zeuzem, Jacques Fellay

**Affiliations:** School of Life Sciences, École Polytechnique Fédérale de Lausanne, Switzerland; Swiss Institute of Bioinformatics, Lausanne, Switzerland; Gilead Sciences Inc, Foster City, CA, United States; Goethe University Hospital, Frankfurt, Germany; Precision Medicine Unit, Lausanne University Hospital, Lausanne, Switzerland

## Abstract

Genetic polymorphism in the interferon lambda (IFN-λ) region is associated with spontaneous clearance of hepatitis C virus (HCV) infection and with response to interferon-based antiviral treatment. Here, we evaluate the associations between IFN- λ polymorphism and HCV variation through a genome-to-genome analysis in 8,729 patients from diverse ancestral backgrounds infected with various HCV genotypes. We searched for associations between rs12979860 genotype, a tag for IFN-λ haplotypes, and amino acid variants in the NS3, NS4A, NS5A and NS5B HCV proteins. We report multiple associations between host and pathogen variants in the full cohort as well as in subgroups defined by viral genotype and human ancestry. We also assess the combined impact of human and HCV variation on pre-treatment viral load. By demonstrating that IFN-λ genetic variation leaves a large footprint in the viral genome, this study provides strong evidence of pervasive viral adaptation to host innate immune pressure during chronic HCV infection.

## Introduction

Infection with hepatitis C virus (HCV), a positive strand RNA virus of the Flaviridae family, represents a major health problem, with an estimated 71 million chronically infected patients worldwide^1^ In the absence of treatment, 15-30% of individuals with chronic HCV infection develop serious complications including cirrhosis, hepatocellular carcinoma and liver failure^2-5^.

Seven major genotypes of HCV have been described, further divided into several subtypes^6,7^ Moreover, within each infected individual, multiple distinct HCV variants co-exist as quasipecies^8^ Inter-host and intra-host HCV evolution is shaped by multiple forces, including human immune pressure^9^ To investigate the complex interactions between host and pathogen at the level of genetic variation, we proposed a genome-to-genome approach that allows the joint analysis of host and pathogen genomic data^10^ Using an unbiased association study framework, a genome-to-genome analysis aims at identifying the escape mutations that accumulate in the pathogen genome in response to host genetic variants. Ansari et al.^11^ used this approach to analyze a cohort of individuals of white ancestry predominantly infected with genotype 3a HCV; they identified associations between viral variants and human polymorphisms in the interferon lambda (IFN-λ) and HLA regions, demonstrating an impact of both innate and acquired immunity on HCV sequence variation during chronic infection.

The IFN-λ association is of particular interest considering the known impact of this polymorphic region on spontaneous clearance of HCV and on response to interferon-based treatment^12-15^ The rs12979860 variant, located 3 kb upstream of the IL28B gene (encoding IFN-λ3), showed the strongest correlation with treatment-induced clearance of infection in the first report^12^ More recent studies have shown that rs12979860 tags a dinucleotide insertion/deletion polymorphism, IFNL4 rs36823481516, which controls generation of the IFN-λ4 protein and is thus the most likely functional variant in the region^28^ The two variants (rs12979860 and rs368234815) are in strong linkage disequilibrium, which explains the differences in HCV outcomes originally associated with rs12979860.

Here, we aim at characterizing the importance of innate immune response in modulating chronic HCV infection by describing the footprint of IFN-λ variation in the viral proteome. Using samples from a heterogeneous cohort of 8,729 HCV-infected individuals, we genotyped the single nucleotide polymorphism (SNP) rs12979860, a known marker of IFN- λ haplotypes, and obtained partial sequences of the HCV genome (*NS3*, *NS4A*, *NS5A* and *NS5B* genes). We tested for associations between rs12979860, HCV amino acid variants and pre-treatment viral load. We show that IFN-λ polymorphism has a pervasive impact on HCV, by describing multiple associations between host and pathogen variants in the full cohort and in subgroups defined by viral genotype or human ancestry. We also present an association analysis of human and viral variants with HCV viral load, which allows for a better understanding of the connections between genomic variation, biological mechanisms and clinical outcomes.

## Results

### Host and pathogen data

We obtained paired human and viral genetic data for 8,729 HCV-infected patients participating in various clinical trials of anti-HCV drugs. The samples were heterogeneous in terms of self-reported ancestry (85% Caucasians, 13% Asians and 2% Africans) and HCV genotypes, with a majority of HCV genotype 1a, 2a and 3a (**Table 1**). On the host side, we genotyped the SNP rs12979860, which reliably tags the known IFN-λ polymorphism in Europeans and Asians^14^ On the pathogen side, we performed deep sequencing of the coding regions of the non-structural proteins NS3, NS4A, NS5A and NS5B^16^ A binary variable was generated for each alternate amino acid, indicating the presence or absence of that allele in a given sample (N = 10,681). For the analysis we used only positions where the amino acid was present in at least 0.3% of the samples (N = 4,022).

**Table 1:**
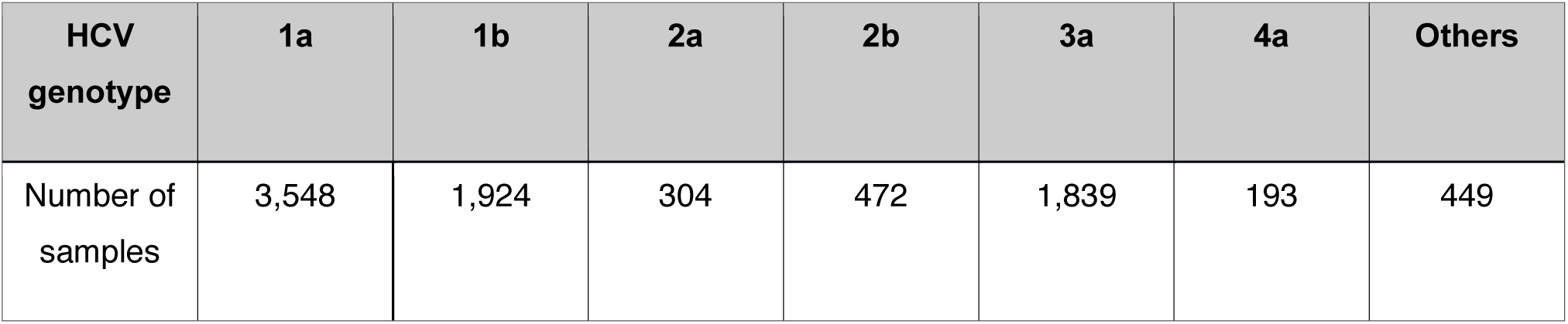
Distribution of HCV genotypes.

| HCV genotype | 1a | 1b | 2a | 2b | 3a | 4a | Others |
| --- | --- | --- | --- | --- | --- | --- | --- |
| Number of samples | 3,548 | 1,924 | 304 | 472 | 1,839 | 193 | 449 |

### Genome-to-genome analyses

We observed highly significant associations between rs12979860 and multiple HCV amino acid variants throughout the HCV proteome (Figure 1). With a Bonferroni correction threshold of 4.7 x 10^−6^ 97 significant association were observed under additive model (**Supplementary Table 1**) and 111 under recessive model (**Supplementary Table 2)**. Presence of Proline at position 156 of NS5B showed the strongest association with rs12979860 genotype (p = 5.2 × 10^−48^ under additive and p = 2.4 × 10^−54^ under recessive model). We observed an enrichment of significant associations (fisher test p = 2 x 10^−4^) in the interferon-sensitivity determining region (ISDR) (aa position 237-276 of NS5A), the strongest one with Lysine at position 265 (p = 1.4 x 10^−12^ under additive and p = 1.3 x 10^−13^ under recessive model). In concordance with a previous study^17^ we also observed a significant association with Histidine at position 93 of NS5A (p = 6.5 x 10^−15^ under additive and p = 2.1 x 10^−17^ under recessive model).

**Figure 1:**
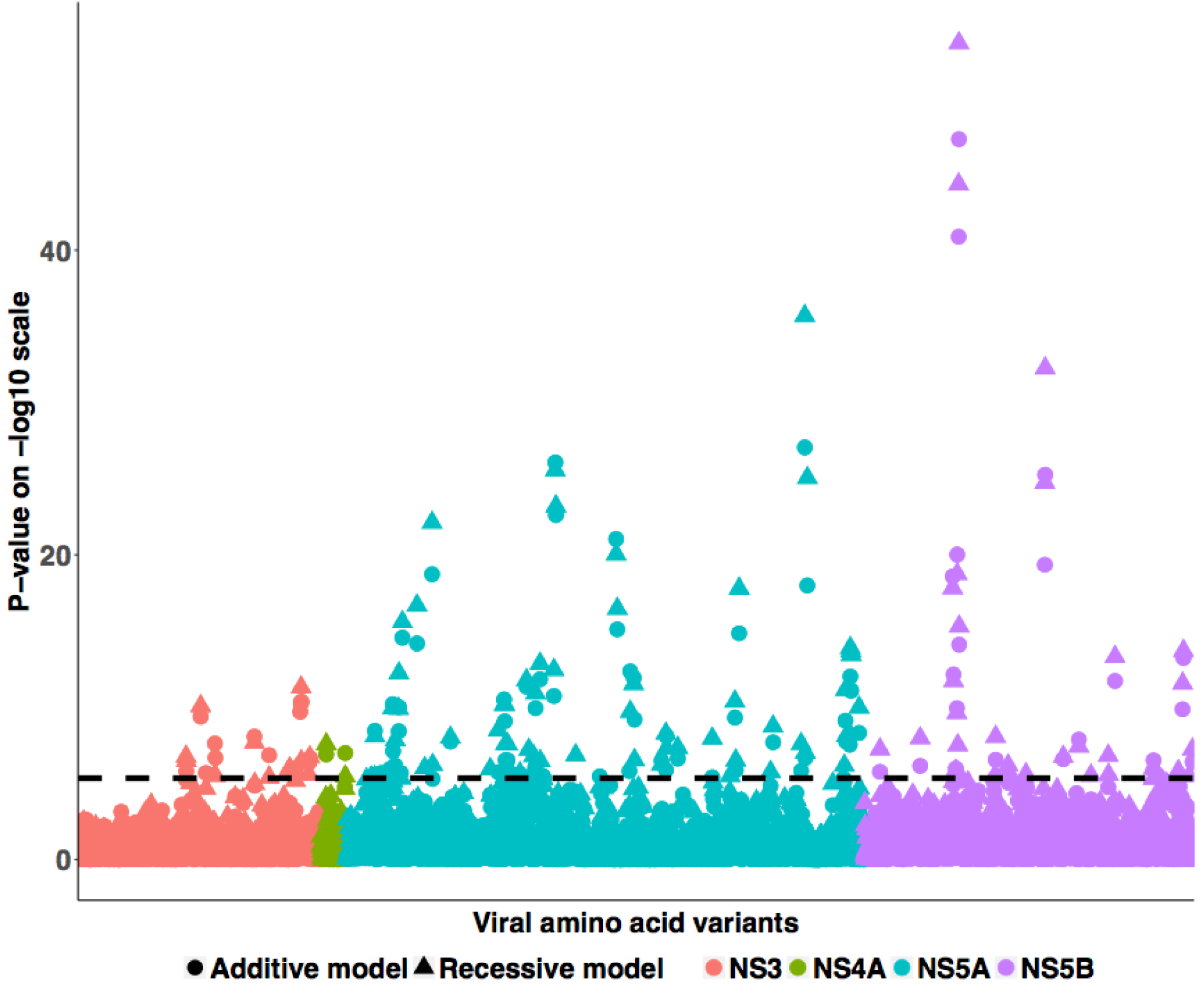
Genome-to-genome analysis results Manhattan plot for associations between human SNP rs12979860 and HCV amino acid variants. The dotted line shows the Bonferroni-corrected significance threshold.

To test for potential heterogeneity of the association signals, we ran sensitivity analyses in various subsets of the study population:

First, we performed separated association studies for each HCV genotype group, restricting the analysis to the following genotypes, present in at least 100 participants: 1a, 1b, 2a, 2b, 3a and 4a. The numbers of significant associations (p_threshold_ < 4.7 x 10^−6^) under additive and recessive models, per genotype, are shown in Figure 2A. The largest number of significant associations was detected for genotype 1a, most likely reflecting an effect of sample size on statistical power. Some associations were significant across genotypes (**Supplementary table 3**). For example, position 156 in NS5B (presence of Proline) was found to be significantly associated with IFN-λ polymorphism in genotypes 1a (additive: p = 4.5 x 10^−6^ recessive: p = 1.2 x 10^−6^), 2b (additive: p = 3.3 x 10^−13^ recessive: p = 2.5 x 10^−13^) and 3a (additive: p = 7.4 x 10^−9^ recessive: p = 1.5 x 10^−9^).

**Figure 2:**
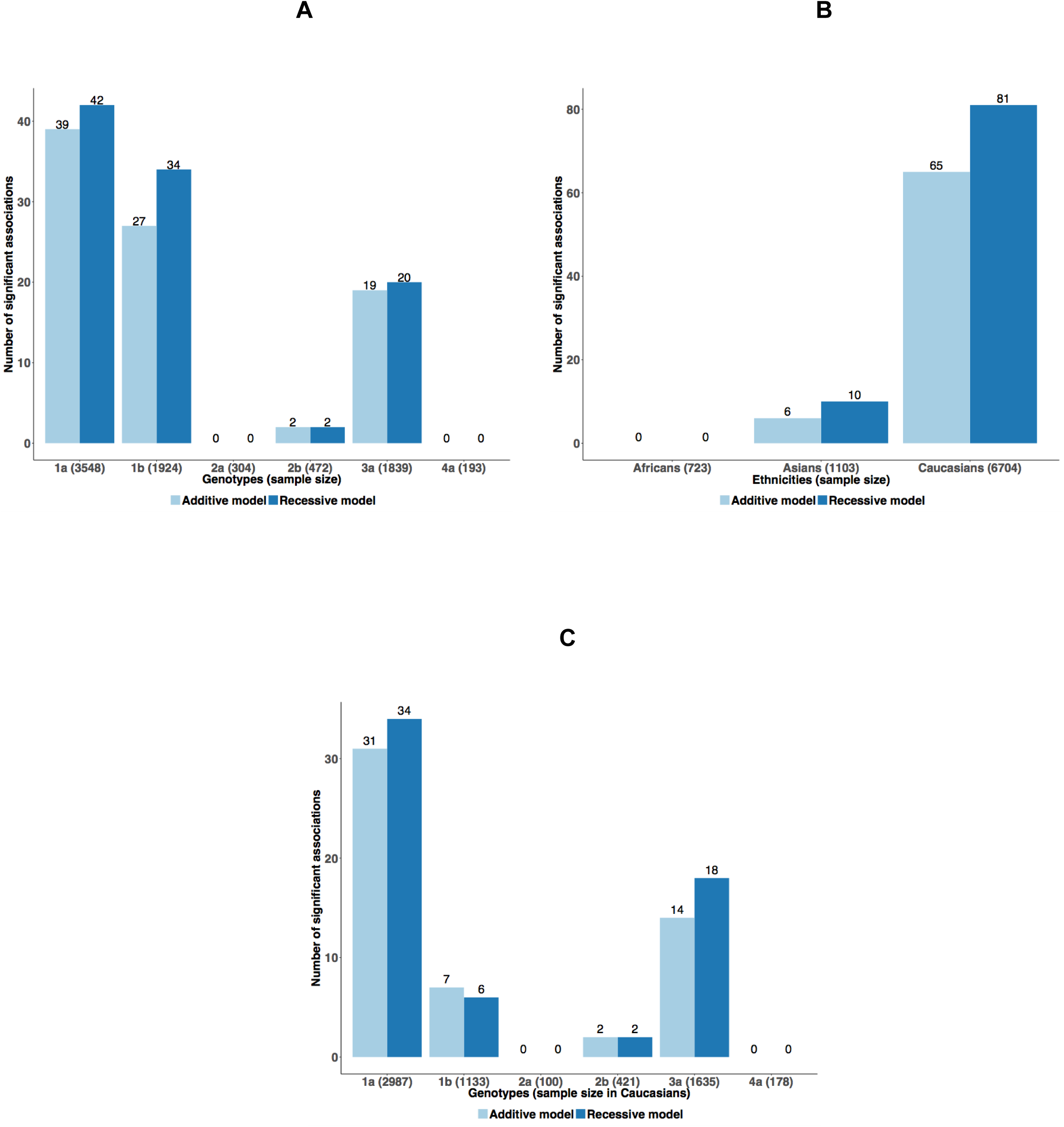
Genome-to-genome analysis results by HCV genotype and ancestry groups (**A**): Bar-plot representing the number of significant genome-to-genome associations per HCV genotype. (**B**): Bar-plot representing the number of significant genome-to-genome associations per self-reported ancestry. (**C**): Bar-plot representing the number of significant genome-to-genome associations per genotype in samples of European ancestry.

Secondly, we ran genome-to-genome analyses in subgroups defined by self-reported ancestry: European, Asian and African. The highest number of significant associations was observed in Europeans, which also represent the largest group (Figure 2B). Here again, presence of Proline at position 156 of NS5B was significantly associated in Asians (additive: p = 3.4 x 10^−6^) and Europeans (additive: p = 2.2 x 10^−40^ recessive: p = 1.9 x 10^−46^) and presence of Histidine at position 93 of NS5A was significantly associated in Europeans (additive: p = 1.7 x 10^−12^ recessive: p = 2.7 x 10^−15^). We also found multiple associations in the ISDR region of NS5A in both Asians and Europeans, including significant association for Lysine at position 265 (**Supplementary table 4)**.

Finally, we further dissected the associations signals within the largest ancestry group, Europeans, by running a per genotype analysis within these samples (Figure 2C). The strongest association was observed for Isoleucine at position 280 of NS5A in patients infected with HCV genotype 1a (additive: p = 3.8 x 10^−24^ recessive: p = 3.3 x 10^−27^). In the same group, we also observed a significant association for Histidine at position 93 of NS5A (additive: p = 2.6 x 10^−22^ recessive: p = 9.8 x 10^−25^). The association with Proline at position 156 of NS5B was not replicated in genotype 1a and 1b samples, but it was found to be significantly associated in the smaller samples of genotype 2b (additive: p = 4.8 x 10^−12^ recessive: p = 2.9 x 10^−12^) and 3a (additive: p = 2.7 x 10^−8^ recessive: p = 6.1 x 10^−9^). All the significant results are presented in **Supplementary table 5.**

### HCV amino acid variants and viral load

To understand the clinical implications of viral mutations associated with IFN-λ polymorphism, we searched for associations between rs12979860, HCV amino acid variants and viral load. Pre-treatment viral load was found to be significantly associated with rs12979860 (p = 2.3 x 10^−16^). In the proteome-wide association study, we identified 112 amino acids significantly associated with viral load (p_threshold_ < 4.7 x 10^−6^) out of 4,022 tested variants. The strongest association was observed for Isoleucine at position 280 of NS5A (p = 1.1 x 10^−30^) (Figure 3A).

**Figure 3:**
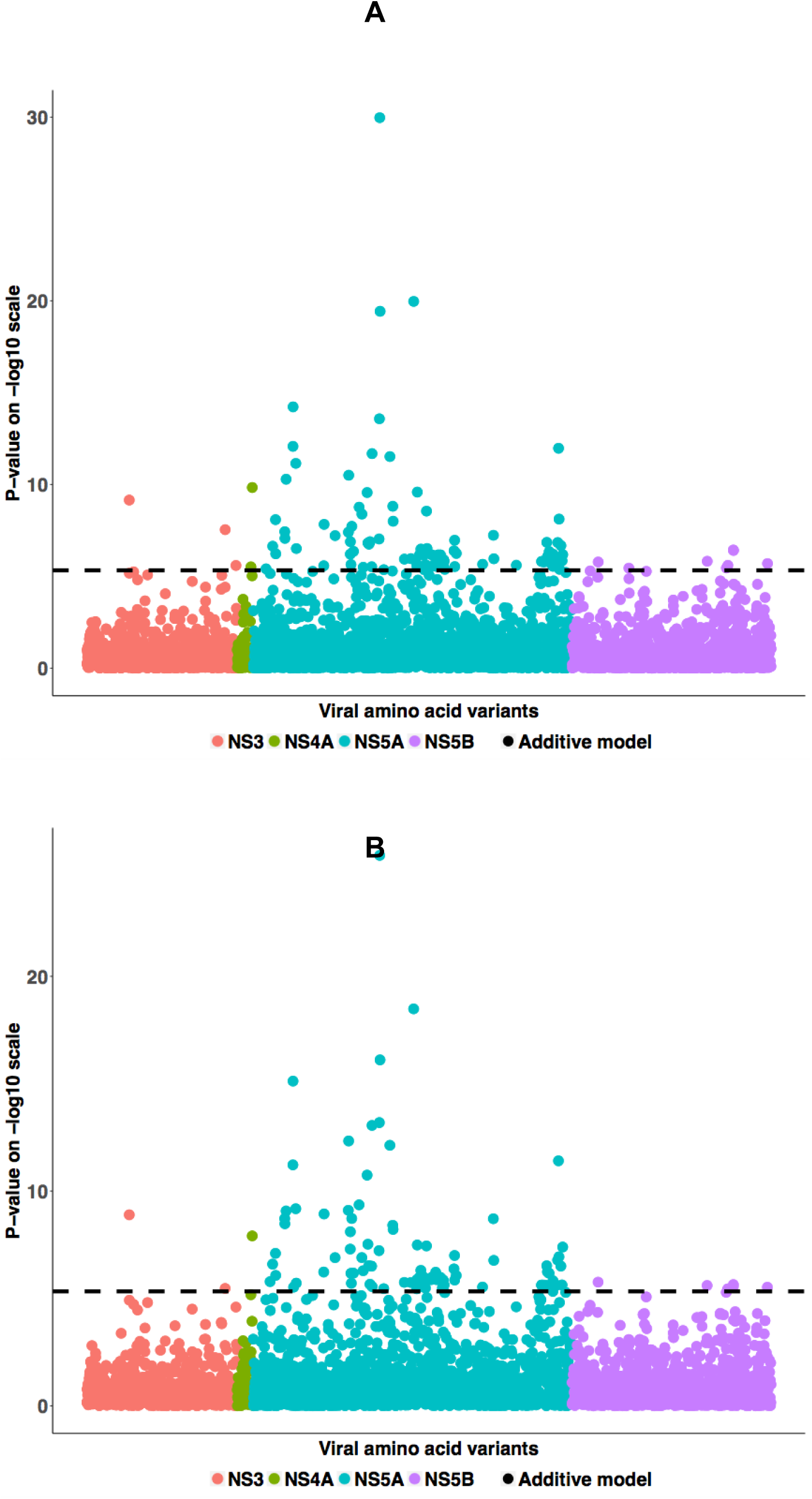
Viral load proteome-wide association results (**A**): Manhattan plot for associations between viral load and HCV amino acid variants. The dotted line shows the Bonferroni-corrected significance threshold. (**B**): Manhattan plot for associations between viral load residuals and HCV amino acid variants.

To further understand these associations, we performed a residual regression analysis. We searched for associations between the HCV amino acid variants and viral load residuals, obtained after regressing viral load on rs12979860. The objective of this analysis was to identify amino acids associated with changes in viral load that cannot be entirely explained by IFN-λ variation. We detected 107 significant associations in total from this analysis (Figure 3B). Interestingly, 22 amino acids, which associated with IFN-λ variations, also showed significant association with viral load as well as viral load residuals (**Supplementary table 6**), including position 265 (presence of Lysine) in the ISDR region of NS5A and position 93 on NS5A (presence of Histidine).

## Discussion

We used an integrated genome-to-genome approach to explore the impact of human genetic variation in the IFN-λ region on the HCV proteome during chronic infection. Our results reveal a strong footprint of innate immune pressure on the non-structural regions of the HCV genome and provide strong evidence for pervasive HCV adaptation to escape human immunity. We also performed sub-analyses in different sample groups, which showed a consistent impact of IFN-λ variation on HCV across genotypes and ancestry categories. Finally, we report viral amino acids significantly associated with both IFN-λ variations and viral load, indicating that some of the HCV clinical and biological outcomes can be explained through host-pathogen interactions.

Our analysis detected multiple associations in all tested proteins, including NS5A. This protein is required for HCV RNA replication and virus assembly and has been shown to associate with interferon signaling and hepatocarcinogenesis^18^ Previous studies have also shown strong associations between variants in the interferon sensitivity determining region of NS5A and viral load as well as response to IFN-based therapy ^19,20^ Some of the strongest associations that we observed were in and around this highly variable interferon sensitivity-determining region of NS5A, suggesting a possible role of these variants in determining the response to IFN-based antiviral treatment. We also detected a strong association for NS5A variant Y93H. This supports the previous findings that reported associations between Y93H mutation, IFN-λ genotype and HCV viral load^17^.

Around 32% of study participants harbored the “favorable” rs12979860 CC genotype, which associates with higher rates of spontaneous viral clearance and of successful response to IFN-based anti-HCV treatment. To measure the impact of the homozygous CC genotype on viral amino acid variants, we also performed association analyses using a recessive model. We detected higher number of significant associations with the recessive model compared to the additive model: patients harboring the CC genotype showed a higher prevalence of amino acid changes, suggesting a stronger immune selective pressure on the virus in homozygous individuals.

This is the first comprehensive analysis of IFN-λ-driven HCV adaptation across different viral genotypes and ancestry groups. In addition to identifying genotype or ancestry specific associations, we observed sites of interaction that were consistent across HCV genotypes and ethnicities; for example, the NS5A variant Y93H. These results indicate that IFN-λ-driven viral adaptation is a part of evolution across HCV genotypes.

In an attempt to delineate the biological impact of these associations, we evaluated the associations between HCV amino acid variants and viral load. We were able to detect a subset of amino acids that associated with both INF-λ variation and HCV viral load, supporting the clinical relevance of host and pathogen interactions. Furthermore, we also performed a similar analysis with residual viral load, i.e. the fraction of the viral load variance that that is not explained by INF-λ variation. We detected a group of viral amino acid variants that associated with SNP variations as well as residual viral load, indicating a stronger role of host-pathogen interactions in explaining the variations in HCV viral load.

Interestingly, only 25% of the host-driven HCV amino acid variants were found to be associated with viral load, indicating that a genome-to-genome analysis can reveal correlations that would go unnoticed in association studies that use more downstream laboratory measurements or clinical outcomes as phenotypes.

IFN-λ polymorphism is the strongest human genetic predictor of spontaneous HCV clearance and response to IFN-based therapy. By integrating INF-λ and HCV amino acid variation in a joint analysis, we here contribute to a better understanding of the genomic mechanisms involved in inter-individual differences in HCV disease outcomes. The large footprint left by INF-λ polymorphism on the HCV proteome is a clear indicator of the central role played by innate immunity in viral control, and of the remarkable capacity of HCV to evolve escape strategies.

## Methods

### Clinical samples

All biological sample were obtained from individuals who provided written informed consent for enrollment in one of 82 clinical studies run by Gilead Sciences (Foster City, CA) and Pharmasset (formerly Princeton, NJ). Study protocols followed the ethical guidelines set in place by the 1975 Declaration of Helsinki and were approved by the relevant institutional review board committees. All samples included in this analysis are baseline samples collected from treatment naive and experienced patients from >25 countries in North America, Europe, Asia, Oceania, and Africa between years 2010 and 2015.

### NS3, NS5A, and NS5B sequencing

The genotype assignment from Siemens VERSANT HCV Genotype INNO-LiPA 2.0 Assay (Innogenetics, Ghent, Belgium) was used to select genotype-specific primers located outside of the gene target(s) that amplify the entire *NS3/4A*, *NS5A*, or *NS5B* regions of HCV. Standard reverse transcription polymerase chain reaction (RT-PCR) was performed on patient plasma with HCV RNA >1000 IU/mL at DDL Diagnostic Laboratory (Rijswijk, The Netherlands). For deep sequencing, amplicons encoding the subject-derived *NS3/4A*, *NS5A* and *NS5B* were run using Illumina MiSeq v2 150 paired-end deep sequencing at DDL or WuXi AppTec (Shanghai, China). FASTQ files were split based on 100% matched barcodes. Contigs were generated from paired-end FASTQ files using VICUNA^21^ and merged to create a de novo assembly sequence. All paired-end reads were merged using PEAR^22^ chopped at the 3’ end when MAPQ<15, and filtered to remove reads <50 bases. The filtered reads were aligned to the de novo assembly sequence using MOSAIK^23^ (v1.1.0017) to create a final assembly sequence. The average coverage of >5000 reads per position was obtained for most of the samples. The aligned reads were translated in-frame and the resulting tabulated summary of variants from the final assembly was utilized to generate a consensus sequence. Mixtures were reported when present in ≥15% of the viral population. NS3/4A, NS5A and NS5B consensus nucleotide and amino acid sequences were compared by the NCBI alignment tool BLAST to a set of reference sequences to assign HCV genotype and subtype. Amino acid variation between the samples that were assigned to genotype 1a, 1b, 2a, 2b, 3a and 4a were tabulated and analyzed. The raw HCV sequences are available in the zenodo repository, https://doi.org/10.5281/zenodo.1476713.

### Host genotyping

Human genotype was determined by means of PCR amplification and sequencing of the rs12979860 single-nucleotide polymorphism, which reliably tags the known IFN-λ polymorphism. Possible genotypes were CC, CT or TT.

### Association analyses

We used logistic regression to search for associations between rs12979860 and the binary pathogen variants, under both additive and recessive models (CC genotype coded as 1), and including sex, country of origin, self-reported ethnicity, cirrhosis status and prior treatment experience as covariates. To account for viral phylogeny, the first 5 phylogenetic principal components^24^ calculated per HCV gene to account for recombination, were also added as covariates. We used muscle^25^ to align the pathogen sequences, RaXML^26^ to obtain the phylogenetic trees and R^27^ for all other analyses.

## Author Contributions

NC and JF designed the analysis methodology. NC performed the analyses and interpretation of data. ES, HM, AO, ML, DB, GS, JM and SZ helped in designing the experiments and acquired the data. JF, NC, ES and AO drafted the manuscript. All authors read and approved the final manuscript.

